# Coding and non-coding drivers of mantle cell lymphoma identified through exome and genome sequencing

**DOI:** 10.1101/686956

**Authors:** Prasath Pararajalingam, Krysta M. Coyle, Sarah E. Arthur, Nicole Thomas, Miguel Alcaide, Barbara Meissner, Merrill Boyle, Bruno M. Grande, Graham Slack, Andrew J. Mungall, Randy D. Gascoyne, Christian Steidl, Joseph Connors, Diego Villa, Marco A. Marra, Nathalie Johnson, David W. Scott, Ryan D. Morin

## Abstract

Mantle cell lymphoma (MCL) is an uncommon B-cell non-Hodgkin lymphoma (NHL) that is incurable with standard therapies. The genetic drivers of this cancer have not been firmly established and the features known to contribute to differences in clinical course remain limited. To extend our understanding of the biological pathways involved in this malignancy, we performed a large-scale genomic analysis of MCL using data from 51 exomes alongside previously published exome cohorts. To confirm our findings, we re-sequenced the genes identified in the exome cohort in 212 MCL tumors, each having clinical follow-up data. We confirmed the prognostic association of *TP53* and *NOTCH1* mutations and further nominate two additional genes, *EWSR1* and *MEF2B*, whose mutation respectively associated with poor and good outcome. Our sequencing revealed novel recurrent mutations including a unique missense hot spot in *MEF2B* and a pattern of non-coding mutations surrounding a single exon of the *HNRNPH1* gene. We sequenced the whole genomes of 34 MCLs to confirm the focal nature of *HNRNPH1* mutations. Using RNA-seq data from 110 of these cases, we identified a functional role for recurrent non-coding *HNRNPH1* mutations in disrupting an auto-regulatory feedback mechanism. Overall, we identified three novel MCL-related genes with roles in RNA trafficking or splicing, namely *DAZAP1, EWSR1*, and *HNRNPH1*. Taken together, these data strongly implicate a role for aberrant regulation of splicing in MCL pathobiology.

**Key points:**

- RNA-binding proteins with roles in regulating alternative splicing, *DAZAP1, EWSR1, HNRNPH1*, are frequently mutated in MCL
- The majority of recurrent somatic *HNRNPH1* mutations are intronic and HNRNPH1 exhibits self-regulation through alternative splicing

## Introduction

Mantle cell lymphoma (MCL) is an uncommon B-cell lymphoma representing 4-9% of non-Hodgkin lymphoma (NHL) diagnoses worldwide^1^. MCL can be broadly divided into two clinical subtypes: nodal and leukemic non-nodal disease, with each displaying distinct natural history, and clinical and genetic features^2^. MCL commonly follows an aggressive clinical course including non-sustained responses to frontline chemo-immunotherapy and frequent relapses, although a subset of cases, including the majority of patients with leukemic non-nodal variant, exhibits significantly longer survival^3^. Clinical prognostic metrics such as MCL International Prognostic Index (MIPI) have enabled patient stratification and improvements to frontline therapy such as the inclusion of active agents (rituximab, bendamustine, cytarabine) as well as consolidative strategies (autologous stem cell transplantation), have significantly improved outcomes over the past two decades^1,4,5^.

The unifying genetic feature of MCL is a chromosomal translocation event that places cyclin D1 *(CCND1)* downstream of the immunoglobulin heavy chain enhancer, causing constitutive *CCND1* expression^2,6^. The translocated *CCND1* allele can also accrue secondary mutations including non-coding mutations in the 3’ untranslated region (UTR), thereby enhancing *CCND1* mRNA stability and further elevating CCND1 protein abundance^7,8^. Through exome and targeted sequencing efforts, largely focused on non-nodal leukemic subtype, several genes have been identified as commonly mutated in MCL, including those involved in DNA damage response *(ATM, TP53)*, epigenetic regulation *(KMT2D, WHSC1)*, Notch signaling *(NOTCH1, NOTCH2)*, NFκB signaling *(CARD11, BIRC3, SYK)*, and ubiquitin mediated proteolysis *(UBR5)*^9–11^. The genomics of MCL have proven to be heterogeneous and diverse, and therefore larger comprehensive explorations are necessary to further understand its biology.

A limited number of recurrent mutations have been associated with prognosis in MCL treated with standard therapy. The most firmly established of these include non-silent mutations affecting *TP53, NOTCH1*, and *CCND1*^12–15^, as well as amplifications of 3q or deletions in 17p^13,16^. With the ongoing evaluation of new therapeutics for MCL, mutations associated with acquired treatment resistance are beginning to be identified^17^. Despite a broad collection of MCL-related genes and mutations, stratification of patients by proliferation, whereby patients are separated into low-, intermediate- and high-risk categories, remains more robust than any individual driver mutation^14,18^. Genetic features that underlie differences in proliferation have yet to be identified.

This study identifies novel recurrent mutations in three RNA-binding proteins *HNRNPH1*, *DAZAP1*, and *EWSR1*, including intronic mutations affecting the splicing of exon 4 in *HNRNPH1*. Our functional characterization in MCL patient samples and cell lines showed *HNRNPH1* splicing is regulated by the HNRNPH1 protein itself via a negative feedback loop leading to alternative inclusion of exon 4 in the mature transcript. By further characterizing the mutational landscape of MCL, we implicate perturbed mRNA processing as an important mechanism in MCL biology.

## Methods

### Study design

We assembled a discovery cohort comprising 51 fresh-frozen diagnostic biopsies from MCL cases collected in British Columbia, Quebec (Montreal), and Ontario and subjected each of these to paired tumor/normal exome sequencing. We included available paired exome data from previous publications to yield a larger discovery cohort of 87 cases. We assembled a validation cohort from FFPE material representing 212 diagnostic tumor samples from BC for targeted sequencing. Only 23 patients overlap between these two cohorts with 189 unique cases in the validation cohort. We also subjected 16 cases from the validation cohort and 18 additional fresh frozen biopsies from BC, along with constitutional DNA, to whole genome sequencing (34 total). We performed RNA-seq on cases spanning all cohorts plus an additional 12 cases from a combination of FFPE and frozen material (110 total). Details for the samples and assays applied to each are included in **Supplemental Data** and **Supplemental Figure 1**. This study was approved by the BC Cancer REB and all participants were recruited with informed consent.

### Exome data analysis

For in-house exomes, we used the Agilent SureSelect Human All Exon kits for library preparation and HiSeq2000 instruments (Illumina) for sequencing. We separately obtained exome data from 29 patients described in *Bea et al.*, which we downloaded from European Genome-Phenome Archive (EGAS00001000510) in BAM format, extracted to FASTQ and validated for data integrity using BamHash (v1.1)^9,19^. FASTQ files for tumor and matched normal exomes (7 patients) described in *Wu et al*. were kindly provided by the authors^11^. External cases qualified for inclusion if a matching normal sample was available. For cases with sequence data from more than one tumor biopsy, we only included the earliest sample in our analysis. We used default settings for all software unless otherwise stated. We used BWA (v0.7.6a) to map reads to the GRCh38 human reference lacking alternate contigs^20^ and subjected these BAM files to soft-clipping of overlapping read pairs using bamUtils clipoverlap (v1.0.13)^21^. For each BAM file, we applied GATK mark duplicates (v3.4.0), and adjusted alignments for putative indels using GATK indel realigner (v3.4.0)^22^. We used Strelka (v1.0.14)^23^ to detect SSMs and indels and annotated these variants using Variant Effect Predictor (Ensembl release 83)^24^ and vcf2maf

### Recurrence analysis

As genes important for lymphomagenesis can exhibit a variety of mutational patterns, we employed a voting strategy involving four separate algorithms to identify recurrently mutated genes and hot spots. Several algorithms to infer genes that are recurrently mutated in cancer cohorts have been described and each of these relies on a variety of features such as predicted functional impact^25,26^, spatial clustering^27^, and mutation rates^28^. We identified significantly mutated genes using a combination of MutSigCV^28^, OncodriveFM^25^, OncodriveFML^26^ and OncodriveCLUST^27^ using a false discovery rate threshold of 0.1 for each algorithm. We promoted genes identified by two or more methods for further sequencing in the validation cohort. Additionally, *NOTCH1, CARD11, NFKBIE* were included due to their importance in MCL and other B-cell NHL^11,12,29^.

### Targeted sequencing, whole genome sequencing and data consolidation

DNA extracted from 212 formalin-fixed paraffin-embedded (FFPE) diagnostic MCL tumor biopsies were used to generate libraries that were enriched for exons corresponding to a set of putative MCL-related genes using a hybridization-based capture approach involving complementary DNA oligonucleotides^30^. These were pooled and sequenced using a MiSeq (Illumina). Paired 150-nt reads were generated, demultiplexed, and mapped to the GRCh38 human reference using Geneious (v9.1.5). Variants with minor allele frequencies greater than 0.0001 in any gnomAD population were considered germline variants and removed^31^. We also performed whole genome sequencing on FF tumor and matched normal samples from 34 MCL cases. Variants from the genome data were identified using Strelka2^32^, which we found to be more robust for identifying variants with low read support or low-level contamination of blood DNA with tumor cells (a feature of some MCLs). We consolidated variants from cases sequenced by more than one method using a tiered approach. Variants found in exomes and genomes were combined per patient. In targeted sequenced tumours for which a normal exome or genome data was available, we considered variants with more than one read support in the normal to be germline variants and removed from analysis. Variants from targeted sequencing cases that did not overlap with exome or genome cases were included in the final variant set. We compared the mutation patterns to diffuse large B-cell lymphoma (DLBCL) using targeted and exome sequencing data from 1616 unique patients^30,33,34^ and removed variants with minor allele frequencies greater than 0.0001 in any gnomAD population^31^.

### RNA-seq analysis of MCL

We performed RNA-sequencing on 110 MCL cases (**Supplemental Methods**). The majority of cases had previously undergone DNA-sequencing, while 12 cases did not. We aligned RNA-seq reads to GRCh38 human reference genome using STAR (version 2.5.3a) followed by Picard MarkDuplicates (version 2.14.1)^22,35^. We used FeatureCounts (version 1.6.0) to separately quantify the reads mapping to the relevant *HNRNPH1* splice junctions using the following parameters: juncCounts, countSplit, ignoreDup, requireBothEndsMapped, and minimum mapping quality ≥ 10. We defined the *HNRNPH1* exon skipping ratio as the ratio of reads spanning the exon 4-6 junction to the sum of reads spanning the exon 4-5 junction and 5-6 junction. RNA-seq samples were designated HNRNPH1 exon 4 mutated or unmutated based on variants discovered in the matching DNA-sequencing sample. In cases where DNA-sequencing data was not available, we determined *HNRNPH1* exon 4 mutation status by examining the exon 4 region in the RNA-sequencing data in IGV.

### Protein-RNA interactions of HNRNPH1

RNA-seq and iCLIP (individual nucleotide resolution cross-linking and immunoprecipitation) data from HeLa cells was provided by authors^36^. Using STAR, we aligned RNA-seq reads from untreated cells transfected with control siRNA (seq8 in Uren data) as described above. Here, we assigned counts using featureCounts with a flattened GTF file containing collapsed transcripts. For HNRNPH iCLIP reads, we pooled reads from two replicates and aligned each pool to GRCh38 with STAR. Peaks in iCLIP data were called by Piranha^37^ in 50 bp bins, accounting for RNA abundance with log-converted RNA-seq counts used as a covariate.

### Cell-based experiments

REC-1 cells were grown as previously described^12,30^. We reconstituted cycloheximide (Sigma) in DMSO and added either cycloheximide or DMSO alone to cells for up to 6 hours. We then extracted RNA or protein, as appropriate, from pellets of 2 × 10^6^ cells using the RNeasy mini kit (Qiagen) or RIPA buffer, respectively, and quantified protein using the Pierce BCA kit (ThermoFisher). Equal amounts of protein were used for SDS-PAGE, and we used Western blotting to observe HNRNPH1 protein levels (abcam ab10374, 1/10 000) and histone H3 protein levels (Cell Signaling Technologies #9715, 1/1 000).

### Digital PCR

For droplet digital PCR (ddPCR), we extracted RNA from FFPE-preserved samples as previously described^18^ and performed reverse transcription with random hexamers (iScript, BioRad) according to manufacturer’s instructions. We pre-amplified cDNA using SsoAdvanced PreAmp SuperMix (BioRad) for 15 cycles with primers targeting *HNRNPH1* (all), *HNRNPH1* (canonical only), *HNRNPH1* (alternative only), *TBP*, *YWHAZ*, and *UBC* (**Supplemental Methods**) as directed. Following a 1 in 5 dilution of pre-amplified cDNA, we performed ddPCR on each on the QX200 system (BioRad) using the primers and probes described in **Supplemental Table 1**. In each sample, we calculated the normalized expression relative to the geometric mean of the expression of three reference genes (*TBP*, *YWHAZ*, and *UBC*). Equal amounts of cell-line RNA were reverse transcribed with random hexamers as above. With diluted cDNA (1 in 10), we performed ddPCR (**Supplemental Table 1**). We normalized expression relative to the geometric mean of three reference genes (*ACTB*, *YWHAZ*, and *UBC*).

### Statistical analysis

All associations between gene mutation status and binary clinical characteristics were assessed using Fisher’s exact test. Association between mutations and overall survival (OS) was determined in the subset of MCL cases that were nodal and treated (n = 175) using the Kaplan-Meier method and log-rank test. We annotated MCL tumours as low-, intermediate- and high-risk based on proliferation gene expression signature using the established Nanostring nCounter-based MCL35 assay^18^. Because the high and intermediate risk cases did not exhibit significantly different overall survival, the intermediate and high proliferation groups were combined into a single group that was defined here as high risk.

## Results

### Somatic Mutation Landscape and Recurrently Mutated Genes

Several genes have previously been implicated as recurrent targets of SSM in MCL^9–11,38^ though many of these candidate drivers have been inconsistent between individual studies^39^. This can be attributed to a combination of genetic heterogeneity and limited cohort sizes. To address this, we sequenced paired tumor/normal exomes from 51 MCLs diagnosed in Canada and analyzed these data alongscide published paired exome data. Three samples exhibited significantly higher mutation burden (median 1761; range 602-8780) and were excluded due to the effect of hypermutation on the detection of drivers, leading to a discovery cohort comprising of 84 cases. In total, 2122 genes were mutated in at least one tumor and tumors, on average, harbored non-silent SSMs in 38 genes (range 11-102).

Through our analysis of this cohort, we found that 16 genes were recurrently mutated as identified by two or more algorithms used to identify driver genes. Three of the algorthims deemed each of *ATM, BIRC3, TP53, S1PR1*, and *B2M* to be significantly mutated, and each of *MEF2B* and *WHSC1* were identified by two methods (**Supplemental Table 2**). Notably, *CCND1*, known to be affected by somatic hypermutation, was identified by OncodriveCLUST, which relies on spatial clustering of mutations. Of the candidate MCL genes, those frequently mutated were *ATM*(44%; n = 37), *CCND1* (18%; n = 15), *TP53* (11%; n = 9), *WHSC1* (15%; n = 13), and *KMT2D* (12%; n = 10), each gene having been previously nominated by other studies (**Supplemental Figure 2**). Three genes not previously attributed to MCL were also identified by at least two methods, including *HNRNPH1* (3.6%; n = 3), *DAZAP1* (3.6%; n = 3), and *EWSR1* (3.6%; n = 3) (**Supplemental Figure 2**). These each encode RNA-binding proteins that play a role in regulating RNA metabolism and alternative splicing of pre-mRNA^40,41^.

### Confirmation of mutation pattern and prevalence

Based on the results and prior studies, we performed targeted sequencing of 18 genes in additional MCLs and used WGS to broadly resolve the exonic and intronic mutation patterns. We applied a combination of targeted sequencing (n = 212) and paired WGS (n = 34) to comprehensively explore the genetic and gene expression landscape of MCL. We consolidated variants across all samples and used the resulting non-duplicated variants for all analyses. Mutation patterns and prevalence in established MCL genes were largely consistent with prior reports (**Figure 1A**; **Supplemental Table 3**). Each of *NOTCH1, MEF2B* and *CCND1* have been shown to have mutation hot spots in MCL and other cancers^7,12,42,43^. In MEF2B, K23R was the predominant mutation observed in MCL (**Supplemental Figure 3**) whereas this mutation was only observed in a single diffuse large B-cell lymphoma (DLBCL) case (n = 1616; P = 4.3 × 10^−17^, Fisher’s exact test). The K23R mutation has been shown to reduce the DNA-binding ability of MEF2B, thus affecting regulation of its transcriptional targets^44^. The majority of *MEF2B* mutations found in DLBCL and follicular lymphomas (FL) affect three hot spots which are absent from MCL, namely K4, D69, and D83^43,45^. These mutations have been described to alter DNA or co-repressor binding by MEF2B, resulting in constitutive expression of MEF2B target genes^46–48^. The distinct hot spot at K23 and paucity of DLBCL/FL-related mutations implies a different role of *MEF2B* mutations in MCL.

**Figure 1.**
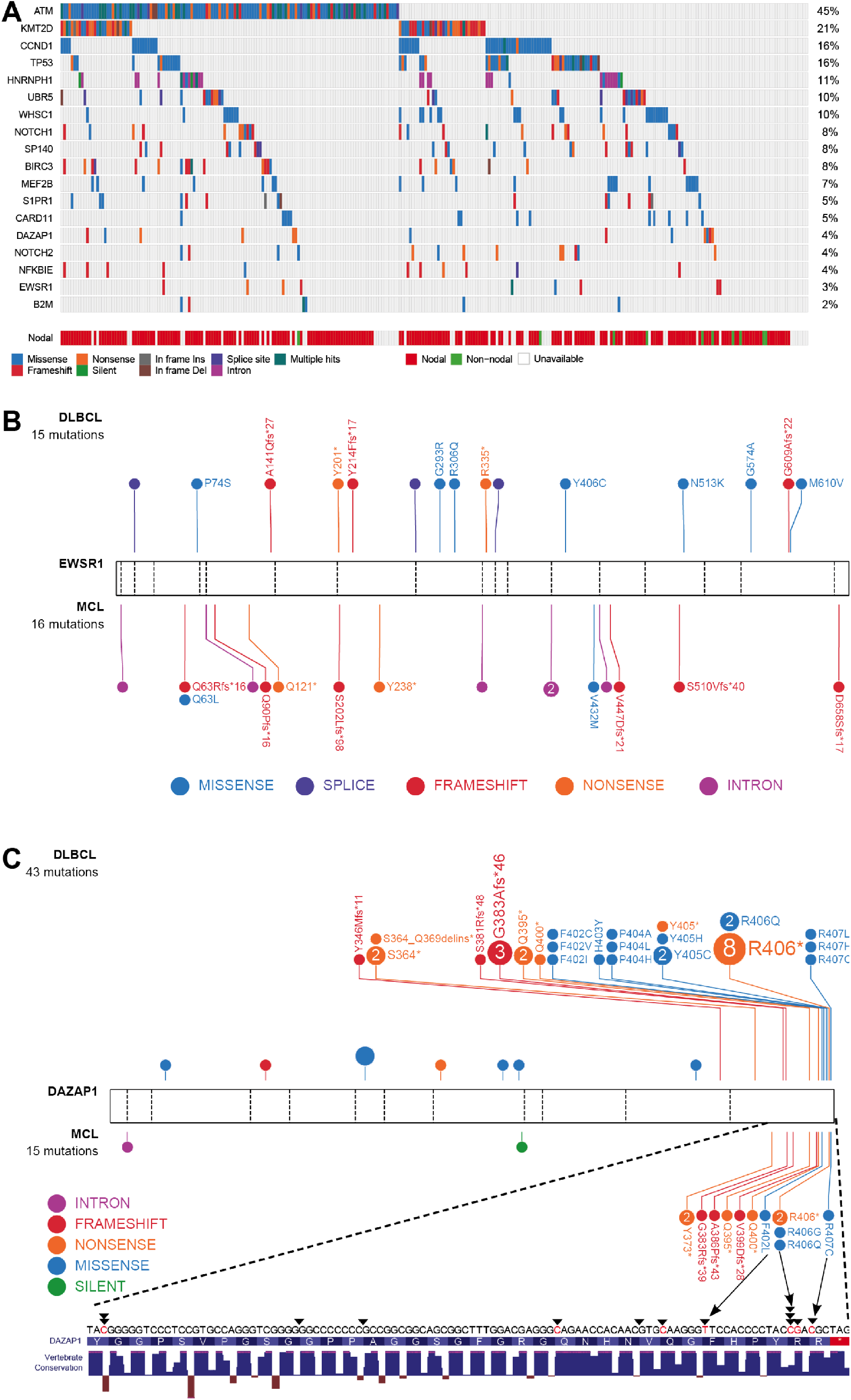
MCL mutation pattern. (A) Mutations observed across 291 MCL samples in 18 candidate MCL genes. Mutations shown here are limited to non-silent mutations for all genes with the exception of *HNRNPH1*, for which intronic and silent mutations were also included. Spatial distribution of mutations observed in (B) *EWSR1*, and (C) *DAZAP1* in MCL (top) compared to DLBCL (bottom).

Unsurprisingly, the incidence of non-silent mutations in newly identified genes was generally lower than those of established MCL genes. *EWSR1* was mutated in 9 cases (3.1%) and DAZAP1, in 13 cases (4.5%) (**Figure 1A**). *EWSR1* predominantly harbored frameshift or nonsense mutations in MCL (n = 8; 2.7%) and exhibited a similar pattern at a lower prevalance in a larger compendium of DLBCLs (n = 5; 0.3%) (**Figure 1B**). This suggests that *EWSR1* has an unappreciated tumor suppressor function in both MCL and DLBCL. *DAZAP1* had a distinctive mutation pattern where mutations were clustered near the C-terminus in a region containing a nuclear localization signal (p.G383-R407)^49^ and proline-rich protein-binding domain^50,51^ (**Figure 1C**). Nine cases harbored putative truncating mutations with each predicted to remove or disrupt the nuclear localization signal while leaving most of the open reading frame intact. Non-synonymous mutations in this region mainly affected highly conserved residues (i.e., p.F402, p.R406, p.R407). Substitution of these residues has been shown to cause cytoplasmic accumulation of DAZAP1 in human kidney epithelial (293T) cells and simian (COS7) cells^49^.

### *HNRNPH1* splice site mutations disrupt HNRNPH1 binding motifs

*HNRNPH1* was mutated in 10% when we considered both coding and non-coding mutations, placing it as the eighth most commonly mutated gene in the cohort (**Supplemental Figure 4**). Despite limited coverage of introns by our sequencing assay, intronic variants were the most common SSM type in this gene, particularly in the regions surrounding exon 4 (**Figure 2A**). Data from paired sequencing confirmed that these were somatic and the WGS data showed they wer largely restricted to this exon and the immediate flanking regions (**Figures 2A and 2B**), HNRNP proteins are widely involved in regulating splicing by binding to pre-mRNA at specific motifs and either promoting or inhibiting usage of nearby splice sites. Distinct from other hnRNPs, HNRNPH1 (and its paralog HNRNPH2) preferentially binds RNA at poly-G motifs^36^. Strikingly, 65% (20/31) of patients with *HNRNPH1* mutations had mutations affecting a poly-G motif within or near this exon. Among the available WGS data from Burkitt lymphoma^52^ (n = 106), DLBCL (n = 153), chronic lymphocytic leukemia (CLL; n = 144) and follicular lymphoma (n = 110), we only identified two DLBCL patients with *HNRNPH1* mutations in this region (1.3%)^30^.

**Figure 2.**
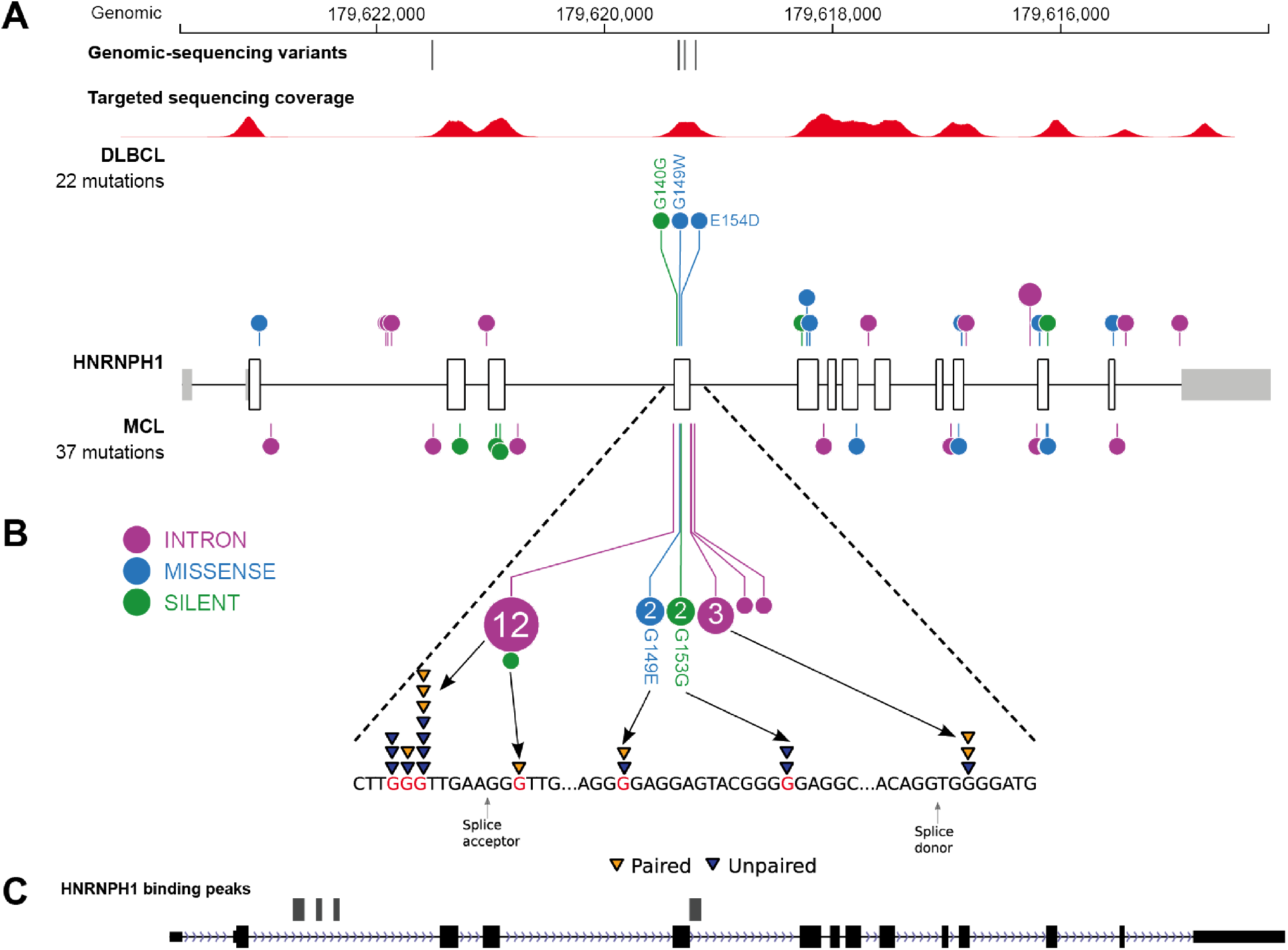
*HNRNPH1* mutations in MCL cluster near exon 4 in poly-G motifs. (A) Somatic mutations found in genomic sequencing cases and targeted sequencing coverage of a representative sample. The prevalence and pattern of mutations in *HNRNPH1* is compared between DLBCL (above) and MCL (below). (B) Splice site and intronic mutations were observed both upstream and downstream of exon 4 affecting poly-G motifs. Paired mutations (orange triangles) are mutations found to be somatic by sequencing matched constitutional DNA (n = 7). Unpaired mutations (blue triangles) are mutations found in tumor-only DNA sequencing (n = 12). (C) HNRNPH1 iCLIP binding peaks show that HNRNPH1 binds near exon 4 of the transcript (shown is Refseq isoform NM_001257293).

The highly specific mutation pattern led us to speculate that HNRNPH1 protein regulates its expression by modulating the splicing of the *HNRNPH1* mRNA transcript. We re-analyzed HNRNPH1 iCLIP-seq data from *Uren et al*.^36^ and confirmed multiple sites of interaction between HNRNPH1 and its pre-mRNA including exon 4 (**Figure 2C**), supporting a model of direct association at these poly-G motifs. *HNRNPH1* has multiple alternative splicing events annotated including the skipping of exon 4. As annotated, isoforms with this event would be subjected to nonsense-mediated decay (NMD) due to disruption of the reading frame. Using a custom ddPCR assay which separately quantifies canonical and alternative *HNRNPH1* transcripts, we demonstrate that inhibition of the NMD process by cycloheximide causes a significant increase in alternative transcript compared to the canonical transcript (**Figure 3A**). This was consistent with a model wherein this gene regulates its own splicing by a negative feedback mechanism to limit HNRNPH1 protein abundance (**Supplemental Figure 5**). Although they do not directly affect canonical splice signals, we hypothesized that the mutations in the poly-G motifs impact the splicing of exon 4.

**Figure 3.**
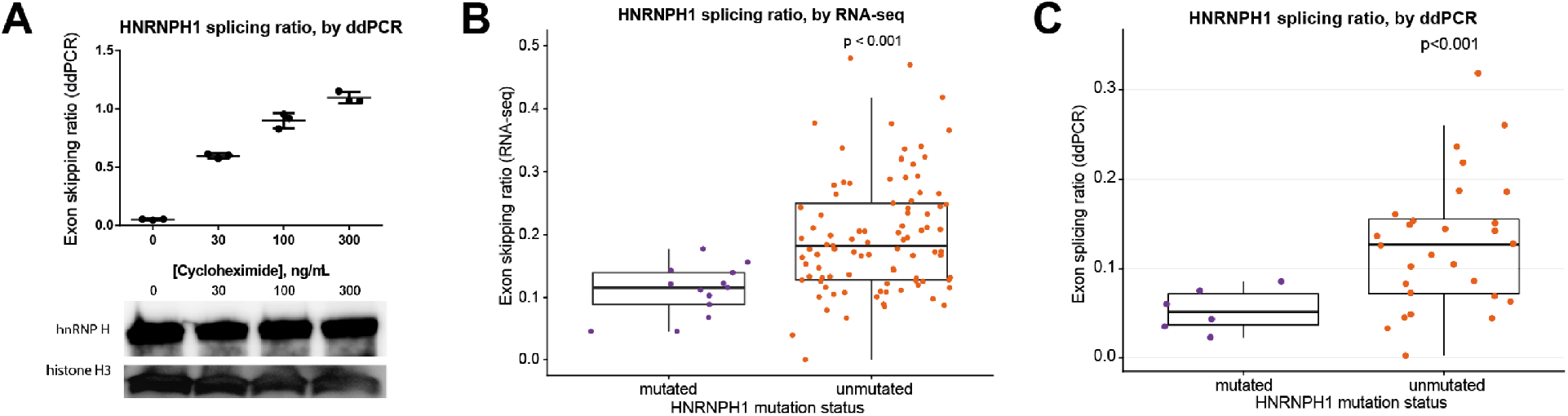
*HNRNPH1* mutations impact RNA splicing. (A) We separately quantified canonical and alternative *HNRNPH1* transcripts by digital PCR in REC-1 cells cultured with cycloheximide (an inhibitor of NMD). This revealed an increasing proportion of the alternative transcript (skipped exon 4) with high concentrations. This increase was not associated with increased HNRNPH protein as determined by immunoblot. (B) Mutated *HNRNPH1* cases showed significantly lower exon skipping ratios compared to unmutated cases, as measured by RNA-seq. (C) Digital PCR was used to separately quantify alternative and canonical *HNRNPH1* transcripts in mutant (n = 6) and wildtype (n = 30) cases. Mutant cases exhibit lower rate of exon skipping and higher overall abundance of HNRNPH1 mRNA.

We analyzed RNA-seq data from 110 cases with known *HNRNPH1* mutation status to evaluate splicing differences between mutated (n = 13) and unmutated (n = 97) tumors. The mutated cases exhibited significantly greater inclusion of exon 4 than the unmutated cases (P = 2.3 × 10^−4^; **Figure 3B**), consistent with our model (**Supplemental Figure 5**). These findings were corroborated by ddPCR of selected cases (P < 0.001; **Figure 3C**), which strongly correlated (R = 0.66, P < 0.01) with splicing ratios identified from RNA-seq data (**Supplemental Figure 5**). These results support the notion that *HNRNPH1* mutations promote the inclusion of exon 4 and the production of mature transcripts by reducing feedback inhibition, most likely by disrupting HNRNPH1 binding motifs surrounding exon 4.

### Association of recurrent mutations with OS

We next explored the relationship between mutation the individual genes and OS in nodal and treated MCL cases that also underwent DNA sequencing. The clinical characteristics of the patients used for survival analysis were largely similar to previous MCL cohorts (**Table 1**). Consistent with previous reports, mutations in *TP53* (**Figure 4A**) and *NOTCH1* (**Figure 4B**) were associated with shorter OS (P = 5.2 × 10 ^(^ and P = 4.0 × 10^−3^, respectively). *MEF2B* K23R mutated cases (n = 9) exhibited significantly longer overall median OS (**Figure 4C**, median not reached; P = 0.04). Among the RNA-binding proteins, we found a novel association between *EWSR1* mutations and worse OS (P = 1.8 × 10^−4^). Given the robust association between proliferation and prognosis, we next examined whether these prognostic mutations were unequally distributed between MCLs with high or low proliferation after combining intermediate risk cases with high risk cases (**Supplemental Figure 6A**). Mutations in either *TP53* (P = 4.3 × 10^−5^) or *MEF2B* (P = 1.4 × 10^−2^) were associated with high-or low-proliferation tumors, respectively (**Supplemental Figure 6B**), while mutations in RNA-processing genes were equally distributed between high- and low-proliferation tumors. Through multivariate analysis we found that only MCL35-derived risk status (P = 1.1 × 10^−2^) and *TP53* mutations (P = 9.6 × 10^−5^) were independently prognostic of OS (**Supplemental Table 5**).

**Figure 4.**
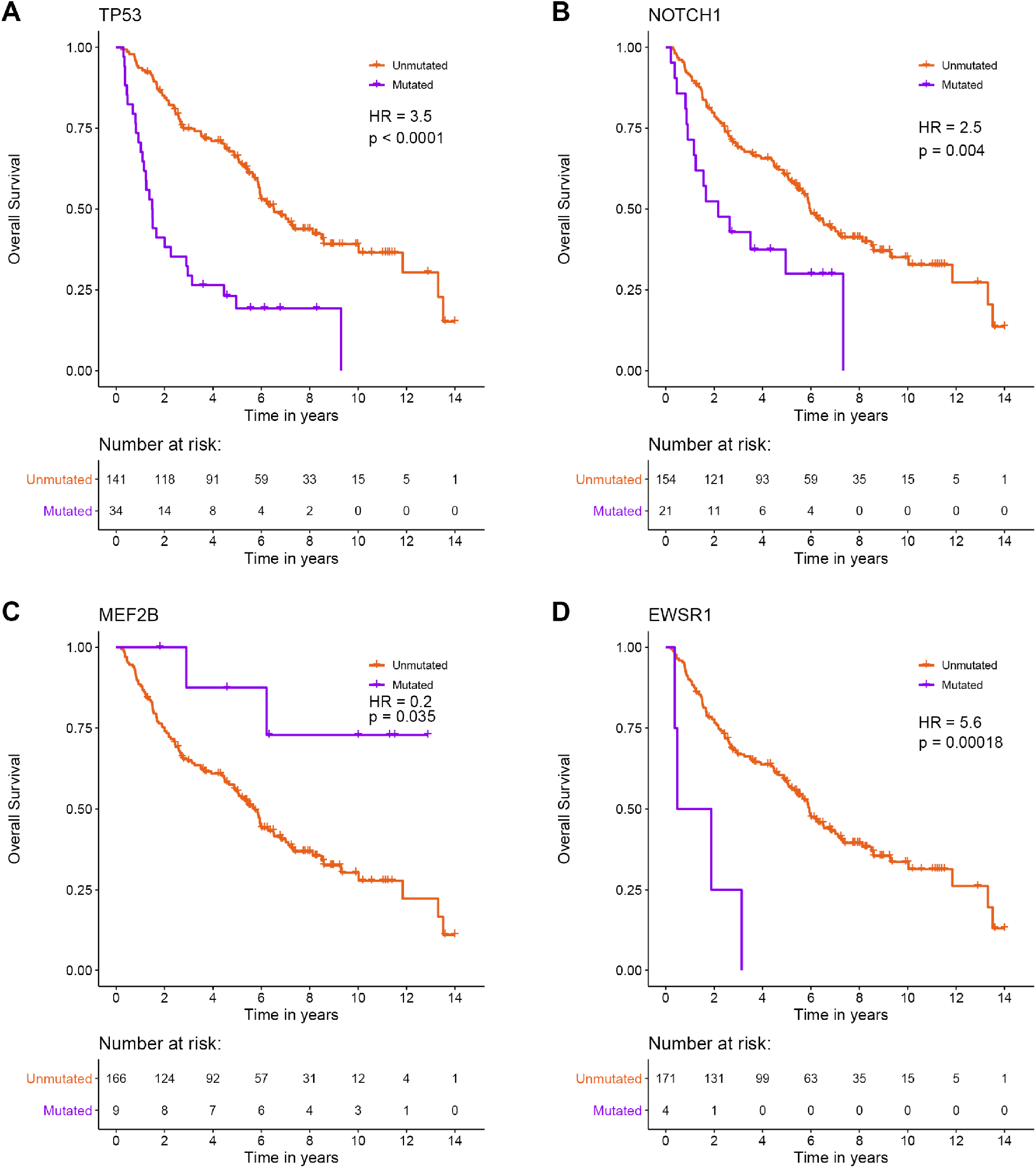
*TP53, NOTCH1, MEF2B*, and *EWSR1* mutations are associated with overall survival in nodal MCL. Survival analysis on the nodal MCL cases revealed four genes with mutations significantly associated with outcome. (A) *TP53* mutated cases (n = 34) had shorter overall survival (P = 5.2 × 10^−9^). (B) *NOTCH1* mutated cases (n = 21) had shorter overall survival (P = 4.0 × 10^−3^). (C) In contrast, *MEF2B* K23R cases (n = 9) exhibited longer overall survival (P = 3.5 × 10^−2^). (D) *EWSR1* mutated cases (n = 4) exhibited shorted overall survival (P = 1.8 × 10^−4^). *TP53, NOTCH1*, and *EWSR1* remained significant after correcting for multiple hypothesis testing (FDR < 0.1).

**Table 1.** Characteristics of patient samples gathered in British Columbia. MCL tumor samples from 232 patients were obtained from patients living in British Columbia. Of the 232 tumors, 175 were nodal and initially treated and were used in survival analysis.

|  | **Nodal, treated (n=175)** | **Others (n=57)\*** | **Total (n=232)** |
| --- | --- | --- | --- |
| **Median age (Range)** | 63 (31-85) | 70 (39-90) | 64 (31-90) |
| **Male (%)** | 153 (77) | 38 (67) | 173 (75) |
| **Performance status >1 (%)** | 37/166 (22) | 8/53 (15) | 45/219 (20) |
| **Elevated LDH (%)** | 54/169 (32) | 4/53 (15) | 58/222 (26) |
| **Blastoid (%)** | 21 (12) | 1 (2) | 22 (10) |
| **Bulky (mass >= 10cm) (%)** | 21 (18) | 5 (9) | 36 (16) |
| **Ann Arbor Stage (%)** |  |  |  |
| 1 | 8/174 (5) | 5/55 (7) | 12/229 (5) |
| 2 | 9/174 (5) | 3/55 (6) | 12/229 (5) |
| 3 | 14/174 (9) | 7/55 (13) | 22/229 (10) |
| 4 | 142/174 (82) | 41/55 (74) | 183/229 (80) |
| **Bone marrow involvement (%)** | 122 (70) | 33 (58) | 155 (67) |
| **GI tract involvement (%)** | 15 (9) | 12 (21) | 27 (12) |
| **B symptoms (%)** | 52/174 (30) | 5/55 (9) | 57/229 (25) |
| **MIPI (%)** |  |  |  |
| High | 28/174 (16) | 2/56 (4) | 30/230 (13) |
| Intermediate | 39/174 (22) | 20/56 (36) | 59/230 (26) |
| Low | 107/174 (62) | 34/56 (61) | 141/230 (61) |
| **Treatment** |  |  |  |
| Splenectomy | 2 | 1 | 3 |
| Palliative chemotherapy | 2 | 0 | 2 |
| CVPR | 3 | 1 | 4 |
| R-CHOP | 153 | 4 | 157 |
| BR | 15 | 1 | 15 |
| Observation | 0 | 51 | 51 |

\* Samples not used in survival analysis: Nodal observed, 47; Non-nodal, 10

## Discussion

This study represents the largest profiling of mutations in MCL to date. Using 291 MCL cases, we validated the recurrence of mutations in genes with known relevance to MCL, including *ATM*, *KMT2D*, *TP53*, *CCND1*, and *NOTCH1*. Using clinical data available for the bulk of these cases, we confirmed the prognostic association of mutations in both *TP53* and *NOTCH1*. *NOTCH1* was not independently prognostic when considering *TP53* mutations and MCL35-derived risk categories. We also found that *WHSC1* mutations were not associated with OS as was previously discovered^53^, which may be due to the limited sample size of that study. We observed prognostic associations for *EWSR1* mutations and *MEF2B* K23R mutations although neither were significant in the multivariate analysis. Confirmation of prognostic association of these genes will require substantially larger cohort sizes. Although the role of *MEF2B* mutations in prognosis remains to be clarified, the high frequency and specificity of K23R mutations in MCL highlights the need for functional characterization of this hot spot as has been done for hot spots common in DLBCL^46,48^.

Althoughly less frequently mutated in MCL, *EWSR1* is an established cancer gene that is typically discussed in the context of the EWSR1-FLI1 fusion oncoprotein that drives Ewing sarcoma^54^. The pattern of mutations observed here implies a separate tumor suppressor role of this gene in MCL and DLBCL. FET family proteins, including EWSR1, each comprise N-terminal transcriptional activating domains and RNA-binding domains near the C-terminus. One of the physiologic functions of EWSR1 entails the coupling of transcription and splicing through its interactions with RNA Polymerase II and recruitment of splicing factors through its C-terminal domain^55–57^. Depletion can induce alternative splicing of EWSR1 targets including genes involved in DNA repair and genotoxic stress signaling^58^. Notably, EWSR1 has been implicated in regulating CCND1 by promoting formation of the less oncogenic CCND1a isoform relative to the shorter CCND1b isoform^59^. Although the targets of EWSR1 have not been established in MCL, our data are consistent with the notion that loss of EWSR1 activity alters RNA metabolism and splicing of genes relevant to MCL.

The DAZAP1 protein contains two N-terminal, highly conserved RNA recognition motif (RRM) domains^60^, which allow it to interact with the pre-mRNA transcripts of its targets and regulate alternative splicing^51,61,62^. Its C-terminal proline-rich domain^63^ is responsible for interactions with other splicing factors^61^. The protein primarily localizes in the nucleus but can also be found in the cytoplasm and within mitochondria^49^. There have been conflicting reports of nuclear localization signals (NLS) in either the N- and C-terminus of this gene^49,64^. A truncated protein lacking the C-terminus or with mutations at highly conserved residues, R393, F402, and F406, has been shown to accumulate in the cytoplasm^49^. In addition to mutations in MCL affecting each of F402 and F406, we observed multiple cases with C-terminal truncating mutations, which would each remove the C-terminal NLS while leaving most of the protein intact. Furthermore, we and others have demonstrated a similar pattern of mutations in a subset of DLBCLs^30,34^. The existence of recurrent *EWSR1* and *DAZAP1* mutations in both malignancies add to the limited genetic features shared between DLBCL and MCL, along with inactivating mutations in *KMT2D* and *TP53*. Based on the effect of mutagenesis experiments^49^, we hypothesize that the commonly observed mutations cause reduced nuclear occupancy of DAZAP1 and affect interactions with other proteins. This could disrupt several processes, including transcription, alternative splicing, mRNA transport and translation^40,51,61^.

HNRNPH1 is a member of the HNRNPH/F family of heterogeneous nuclear ribonucleoproteins^65^. HNRNPH1 binds to various cis-regulatory elements, and depending on the sequence context and interacting proteins, can promote or suppress the use of nearby splice sites^66^. Our data support a model wherein HNRNPH1 protein normally limits its own accumulation by suppressing the inclusion of exon 4 and directing its mRNA for degradation by NMD. In this model, the observed intronic mutations would destroy motifs surrounding exon 4 thereby favoring exon inclusion and enhancing the production of the mature transcript, an observation that is supported by RNA-seq (110 cases) and ddPCR (34 cases) analysis. The possibility that exon 4 inclusion is correlated with HNRNPH1 levels, and the paucity of mutations in this region in other B-cell NHL may indicate a differential role for HNRNPH1 in B-cell development or lymphomagenesis and are consistent with a more important role of HNRNPH1 in MCL biology. Similar to the predicted effects of other RNA binding proteins with a multiplicity of targets, enhanced activity of HNRNPH1 is expected to have widespread effects on the splicing landscape in MCL^36^.

Mutations in these novel MCL-related genes (*EWSR1*, *DAZAP1*, and *HNRNPH1*) are compelling as they implicate mRNA maturation, splicing and/or trafficking in lymphomagenesis. Evidence is accumulating which relates alterations in RNA-binding proteins and splice factors in numerous cancers to various aspects of cancer cell biology. Specifically, small changes in RNA-binding proteins can have large downstream effects on gene expression and can thus impact multiple hallmarks of cancer^67^. For example, the splicing factor SF3B1 was identified as recurrently mutated in CLL^68–70^ and further detailed investigations have identified widespread alternative splicing affecting multiple cellular pathways^71,72^. The identification of pleiotropic downstream effects including DNA damage response, apoptosis, and Notch signaling^71,73^, indicate that widespread disruptions to RNA processing can enhance cancer cell survival by multiple pathways. The evidence for alterations affecting multiple splicing factors in the literature^74,75^, including those observed in this study, suggest an emerging role of RNA metabolism and splicing in B-cell lymphomas.

In summary, through genomic analysis of 303 MCL tumors, we identified novel recurrently mutated genes with a range of mutation incidences. We implicate an important role for RNA-binding proteins and RNA processing in MCL as compared to other B-cell lymphomas, suggesting that RNA metabolism and splicing have a specific role in MCL pathology, although the downstream targets of these genes in MCL have yet to be characterized. Further work which links these mutations to dysregulation of specific RNA molecules will highlight the relevance of RNA processing in MCL.

## Acknowledgments

We thank Canada's Michael Smith Genome Sciences Centre for library construction, sequencing and bioinformatics support and thank Arezoo Mohajeri for the construction of exome libraries and thank Arezoo Mohajeri for the construction of exome libraries. This work was supported by the Canadian Institutes of Health Research (300738), the Terry Fox Research Institute (1021, 1043 and 1061). Biological materials were provided by the Ontario Tumour Bank, which is supported by the Ontario Institute for Cancer Research through funding provided by the Government of Ontario. KMC is supported by a CIHR postdoctoral fellowship.

## Author Contributions

PP analyzed and interpreted the data; SEA, MA performed library preparation; KMC and NT analyzed CLIP-seq and RNA-sequencing data; KMC performed cell-based and FFPE-based assays; MA performed sequencing and analysis targeted sequencing; BMG provided bioinformatic support and interpreted the data; BM, MB, GS, AM performed nucleic acid extractions and sample quality control; DV and DWS provided clinical data and reviewed the cases; DWS, MAM, NJ, RG, CS, JC and RDM interpreted data, designed the study, and with PP, wrote the manuscript.

