## Supplemental Document for "Coding and non-coding drivers of mantle cell lymphoma identified through exome and genome sequencing"

### Data

#### Description of Additional Supplementary Files

File Name: **Supplementary Data 1**

Description: Sample Overview. Metadata for each patient (including patients obtained from public repositories), the sequencing method used, risk category (“MCL35cat”), biopsy site, initial treatment regimen, histopathology and treatment category (“Category”).

#### Methods

**Whole genome sequencing library construction and sequencing**

Whole genome sequencing (WGS) libraries were constructed from MCL fresh frozen tumour and constitutive DNA collected from patients in British Columbia, Canada. To minimize library bias and coverage gaps associated with PCR amplification of high GC or AT-rich regions we have implemented a version of the TruSeq DNA PCR-free kit (E6875-6877B-GSC, New England Biolabs), automated on a Microlab NIMBUS liquid handling robot (Hamilton). Briefly, 500 ng of genomic DNA was arrayed in a 96-well microtitre plate and subjected to shearing by sonication (Covaris LE220). Sheared DNA was end-repaired and size selected using paramagnetic PCRClean DX beads (C-1003-450, Aline Biosciences) targeting a 300-400 bp fraction. After 3' A-tailing, full length TruSeq adapters were ligated. Libraries were purified using paramagnetic (Aline Biosciences) beads. PCR-free genome library concentrations were quantified using a qPCR Library Quantification kit (KAPA, KK4824) prior to sequencing with paired-end 150 nucleotide reads on the Illumina HiSeqX platform using V4 chemistry according to manufacturer recommendations.

**Ribosomal RNA depletion RNA sequencing library construction and sequencing**

To remove cytoplasmic and mitochondrial ribosomal RNA (rRNA) species from total RNA NEBNext rRNA Depletion Kit for Human/Mouse/Rat was used (NEB, E6310X). Enzymatic reactions were set-up in a 96-well plate (Thermo Fisher Scientific) on a Microlab NIMBUS liquid handler (Hamilton Robotics, USA). 100ng of DNase I treated total RNA in 6 µL was hybridized to rRNA probes in a 7.5 µL reaction. Heat-sealed plates were incubated at 95˚C for 2 minutes followed by incremental reduction in temperature by 0.1˚C per second to 22˚C (730 cycles). The rRNA in DNA hybrids were digested using RNase H in a 10 µL reaction incubated in a thermocycler at 37˚C for 30 minutes. To remove excess rRNA probes (DNA) and residual genomic DNA contamination, DNase I was added in a total reaction volume of 25 µL and incubated at 37˚C for 30 minutes. RNA was purified using RNA MagClean DX beads (Aline Biosciences, USA) with 15 minutes of binding time, 7 minutes clearing on a magnet followed by two 70% ethanol washes, 5 minutes to air dry the RNA pellet and elution in 36 µL DEPC water. The plate containing RNA was stored at -80˚C prior to cDNA synthesis.

First-strand cDNA was synthesized from the purified RNA (minus rRNA) using the Maxima H Minus First Strand cDNA Synthesis kit (Thermo-Fisher, USA) and random hexamer primers at a concentration of 8 ng/µL along with a final concentration of 0.4 µg/µL Actinomycin D, followed by PCR Clean DX bead purification on a Microlab NIMBUS robot (Hamilton Robotics, USA). The second strand cDNA was synthesized following the NEBNext Ultra Directional Second Strand cDNA Synthesis protocol (NEB) that incorporates dUTP in the dNTP mix, allowing the second strand to be digested using USERTM enzyme (NEB) in the post-adapter ligation reaction and thus achieving strand specificity.

cDNA was fragmented by Covaris LE220 sonication for 130 s (2x65 s) at a "Duty cycle" of 30%, 450 Peak Incident Power (W) and 200 Cycles per Burst in a 96-well microTUBE Plate (P/N: 520078) to achieve 200-250 bp average fragment lengths. The paired-end sequencing library was prepared following the BC Cancer Agency Genome Sciences Centre strand-specific, plate-based library construction protocol on a Microlab NIMBUS robot (Hamilton Robotics, USA). Briefly, the sheared cDNA was subject to end-repair and phosphorylation in a single reaction using an enzyme premix (NEB) containing T4 DNA polymerase, Klenow DNA Polymerase and T4 polynucleotide kinase, incubated at 20˚C for 30 minutes. Repaired cDNA was purified in 96-well format using PCR Clean DX beads (Aline Biosciences, USA), and 3' Atailed (adenylation) using Klenow fragment (3' to 5' exo minus) and incubation at 37˚C for 30 minutes prior to enzyme heat inactivation. Illumina PE adapters were ligated at 20˚C for 15 minutes. The adapter-ligated products were purified using PCR Clean DX beads, then digested with USERTM enzyme (1 U/µL, NEB) at 37˚C for 15 minutes followed immediately by 13 cycles of indexed PCR using Phusion DNA Polymerase (Thermo Fisher Scientific Inc. USA) and Illumina's PE primer set. PCR parameters: 98˚C for 1 minute followed by 13 cycles of 98˚C 15 seconds, 65˚C 30 seconds and 72˚C 30 seconds, and then 72˚C 5 minutes. The PCR products were purified and size-selected using a 1:1 PCR Clean DX beads-to-sample ratio (twice), and the eluted DNA quality was assessed with Caliper LabChip GX for DNA samples using the High Sensitivity Assay (PerkinElmer, Inc. USA) and quantified using a Quant-iT dsDNA High Sensitivity Assay Kit on a Qubit fluorometer (Invitrogen) prior to library pooling and size-corrected final molar concentration calculation for Illumina HiSeq2500 sequencing with paired-end 75 base reads.

### Tables

**Supplemental Table 1. Primers and probes used for preamplification and digital PCR.**

| Target | Oligo | Sequence |
|---|---|---|
| HNRNPH1 (all) | Forward | TGAAATCTTTAAGAGCAGTAGAG |
| | Reverse | CCCCAGGTCTGTCATAAG |
| | Probe | TCATTATGATCCACCACGA (FAM) |
| HNRNPH1 canonical transcript | Forward | CGGCTTAGAGGACTTCC |
| | Reverse | CACCGGCAATGTTATCC |
| | Probe | TTCTTCTCAGGGTTGGAAAT (FAM) |
| HNRNPH1 alternative transcript | Forward | CGGCTTAGAGGACTTCC |
| | Reverse | AGCTTTCGTGGTGGATC |
| | Probe | CAGTTCTTCTCAGGTATATTGA (HEX) |
| TBP | Forward | GGAGCTGTGATGTGAAGTT |
| | Reverse | AGGAAATAACTCTGGCTCAT |
| | Probe | TAGAAGGCCTTGTGCTCACC (FAM) |
| YWHAZ | Forward | AGAAAGCCTGCTCTCTTG |
| | Reverse | CGTGCTGTCTTTGTATGACT |
| | Probe | TGAAGCCATTGCTGAACTTG (FAM) |
| UBC | Forward | AGACACTCACTGGCAAGA |
| | Reverse | TTTGCTTTGACGTTCTCG |
| | Probe | ATCACCCTTGAGGTCGAGC (FAM) |
| ACTB | Forward | ATGCAGAAGGAGATCACTG |
| | Reverse | AGTACTTGCGCTCAGGA |
| | Probe | CCCAGCACAATGAAGATCAA (FAM) |

**Supplemental Table 2. Recurrently mutated genes in MCL exomes**. Four methods were applied to determine recurrently mutated genes. Several genes were identified by more than one approach. Numbers of mutated cases is based on the discovery cohort of 84 MCL exomes.

| Gene | # Tools | Mutated cases (%) | MutSigCV | OncodriveFML | OncodriveFM | OncodriveCLUST |
|---|---|---|---|---|---|---|
| EWSR1 | 4 | 3 (3.6) | 1 | 1 | 1 | 1 |
| ATM | 3 | 37 (44) | 1 | 1 | 1 | 0 |
| B2M | 3 | 3 (3.6) | 1 | 0 | 1 | 1 |
| BIRC3 | 3 | 8 (9.5) | 1 | 1 | 1 | 0 |
| DAZAP1 | 3 | 3 (3.6) | 1 | 1 | 1 | 0 |
| KMT2D | 3 | 10 (12) | 1 | 1 | 1 | 0 |
| NOTCH2 | 3 | 4 (4.8) | 0 | 1 | 1 | 1 |
| S1PR1 | 3 | 6 (7.1) | 1 | 1 | 1 | 0 |
| TP53 | 3 | 9 (11) | 1 | 1 | 1 | 0 |
| UBR5 | 3 | 6 (7.1) | 1 | 1 | 1 | 0 |
| GPR32 | 2 | 4 (4.8) | 0 | 0 | 1 | 1 |
| HNRNPH1 | 2 | 3 (3.6) | 1 | 0 | 1 | 0 |
| MEF2B | 2 | 8 (9.5) | 1 | 0 | 1 | 0 |
| SP140 | 2 | 5 (6) | 1 | 0 | 1 | 0 |
| SPEN | 2 | 3 (3.6) | 0 | 1 | 1 | 0 |
| WHSC1 | 2 | 13 (15) | 0 | 1 | 1 | 0 |

**Supplemental Table 3. Mutation frequency in MCL genes**. Mutation counts of panel genes. “Paired Total” is the number of mutations found in tumour-normal paired analyses (i.e., exomes and/or genomes). Abbreviations: UTR, untranslated region.

| Gene | Flank | Frameshift | In frame Del | In frame Ins | Intron | Missense | Nonsense | Silent | Splice site | UTR | Total Mutations | Paired Total |
|---|---|---|---|---|---|---|---|---|---|---|---|---|
| CCND1 | 12 | 0 | 0 | 0 | 74 | 58 | 2 | 44 | 0 | 33 | 223 | 44 |
| ATM | 0 | 21 | 4 | 0 | 13 | 107 | 31 | 3 | 15 | 0 | 194 | 60 |
| KMT2D | 1 | 22 | 2 | 1 | 3 | 22 | 21 | 10 | 2 | 5 | 89 | 23 |
| TP53 | 1 | 3 | 2 | 0 | 7 | 33 | 12 | 0 | 2 | 0 | 60 | 18 |
| UBR5 | 1 | 7 | 1 | 0 | 13 | 13 | 4 | 2 | 9 | 2 | 52 | 21 |
| WHSC1 | 0 | 0 | 0 | 0 | 16 | 32 | 0 | 1 | 0 | 1 | 50 | 16 |
| NOTCH1 | 3 | 11 | 0 | 0 | 6 | 8 | 7 | 5 | 1 | 2 | 43 | 12 |
| HNRNPH1 | 0 | 0 | 0 | 0 | 25 | 6 | 0 | 6 | 0 | 2 | 39 | 12 |
| SP140 | 0 | 8 | 0 | 0 | 11 | 11 | 1 | 1 | 3 | 0 | 35 | 13 |
| NOTCH2 | 0 | 2 | 0 | 0 | 12 | 6 | 6 | 5 | 0 | 3 | 34 | 12 |
| CARD11 | 3 | 0 | 0 | 0 | 6 | 15 | 0 | 1 | 0 | 4 | 29 | 12 |
| BIRC3 | 0 | 12 | 1 | 0 | 4 | 3 | 7 | 1 | 1 | 0 | 29 | 16 |
| MEF2B | 0 | 0 | 0 | 0 | 5 | 22 | 0 | 0 | 0 | 0 | 27 | 13 |
| S1PR1 | 2 | 6 | 1 | 2 | 0 | 7 | 0 | 0 | 0 | 0 | 18 | 12 |
| DAZAP1 | 1 | 3 | 0 | 0 | 1 | 4 | 6 | 1 | 0 | 2 | 18 | 5 |
| NFKBIE | 0 | 9 | 0 | 0 | 3 | 1 | 1 | 2 | 1 | 0 | 17 | 3 |
| EWSR1 | 0 | 6 | 0 | 0 | 6 | 2 | 2 | 0 | 0 | 0 | 16 | 8 |
| B2M | 0 | 2 | 0 | 0 | 0 | 4 | 1 | 0 | 0 | 1 | 8 | 5 |

**Supplemental Table 4. HNRNPH1 mutations identified in all MCL cases sequenced**

| Chromosome | Start | End | Variant Type | Sample | Reference | Alternative |
|---|---|---|---|---|---|---|
| chr5 | 179619260 | 179619268 | Intron | 10-40390 | C | T |
| chr5 | 179619260 | 179619268 | Intron | P6 | C | T |
| chr5 | 179619260 | 179619268 | Intron | 10-30161 | C | T |
| chr5 | 179619340 | 179619362 | Silent | 05-22043 | C | G |
| chr5 | 179619340 | 179619362 | Silent | 08-29218 | C | T |
| chr5 | 179619340 | 179619362 | Missense | 15-30071 | C | T |
| chr5 | 179619340 | 179619362 | Missense | 09-17287 | C | T |
| chr5 | 179619405 | 179619420 | Missense | 16-32881 | C | A |
| chr5 | 179619405 | 179619420 | Intron | 01-15563 | C | A |
| chr5 | 179619405 | 179619420 | Intron | 16-31255 | C | A |
| chr5 | 179619405 | 179619420 | Intron | 13-14456 | C | A |
| chr5 | 179619405 | 179619420 | Intron | 02-12941 | C | A |
| chr5 | 179619405 | 179619420 | Intron | 08-36779 | C | A |
| chr5 | 179619405 | 179619420 | Intron | 12-23793 | C | A |
| chr5 | 179619405 | 179619420 | Intron | 96-24127 | C | A |
| chr5 | 179619405 | 179619420 | Intron | M022 | C | G |
| chr5 | 179619405 | 179619420 | Intron | 05-14198 | C | G |
| chr5 | 179619405 | 179619420 | Intron | 07-30311 | C | A |
| chr5 | 179619405 | 179619420 | Intron | 09-16272 | C | A |
| chr5 | 179619405 | 179619420 | Intron | 95-32777 | C | A |

**Supplemental Table 5. Multivariate survival analysis summary**. Hazard ratios for mutated vs. unmutated *TP53*, *EWSR1*, *NOTCH1*, and *MEF2B*, and low- vs. high-risk category were calculated for nodal, treated cases (n = 175).

| Covariate | Hazard ratio | Std. error | Statistic | P-value | CI low | CI high |
|---|---|---|---|---|---|---|
| TP53 | 0.99 | 0.25 | 3.9 | $9.6 \times 10^{-5}$ | 0.49 | 1.5 |
| MCL35catLow | -0.59 | 0.23 | -2.5 | 0.01 | -1.05 | -0.14 |
| NOTCH1 | 0.54 | 0.30 | 1.8 | 0.07 | -0.04 | 1.14 |
| EWSR1 | 0.98 | 0.54 | 1.8 | 0.07 | -0.09 | 2.05 |
| MEF2B | -17.2 | 2902.4 | -0.006 | 0.99 | -Inf | Inf |

Abbreviations: CI, 95% confidence intervals.

### Figures

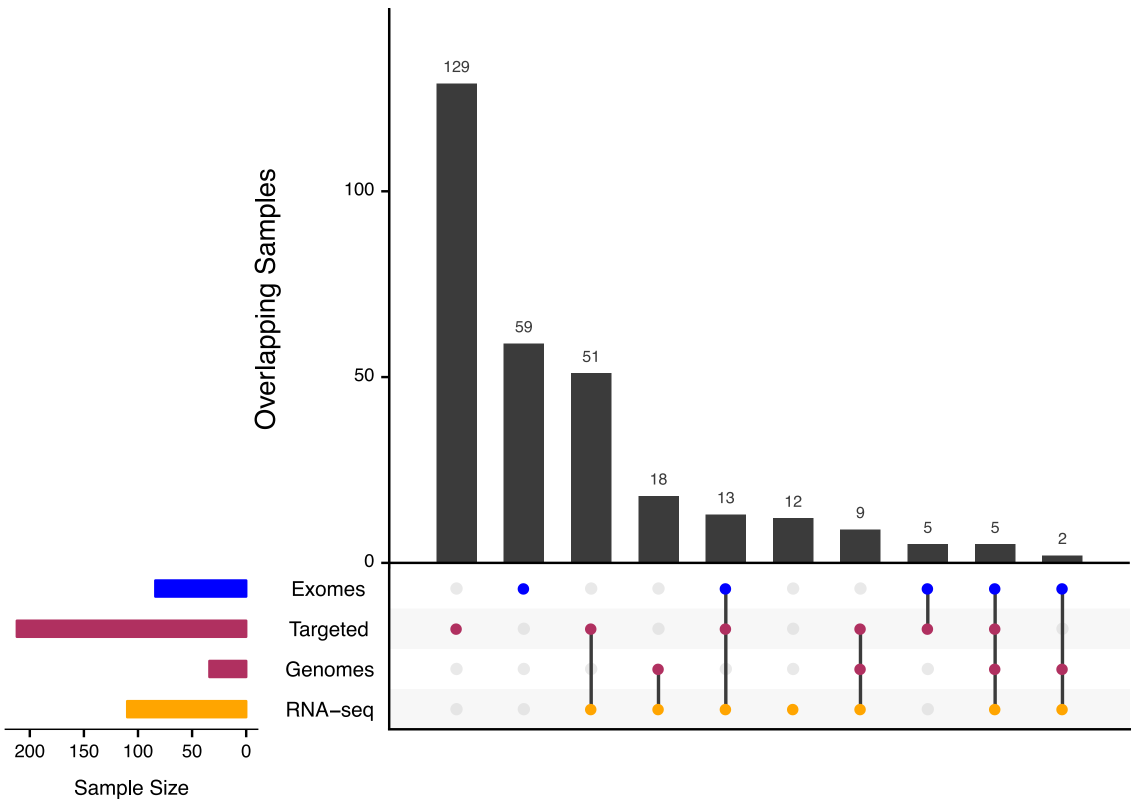


**Supplemental Figure 1. Overlapping samples used mutation discovery and validation.** Data sets are colored according its inclusion during the study: blue, discovery cohort; maroon, validation cohort; orange, *HNRNPH1* splicing validation. Data consolidation began by merging variants from exomes and genomes. For targeted sequencing data, in cases where a normal exome or genome was available, variants were pooled and those with more than one read support in the normal were considered germline and removed.

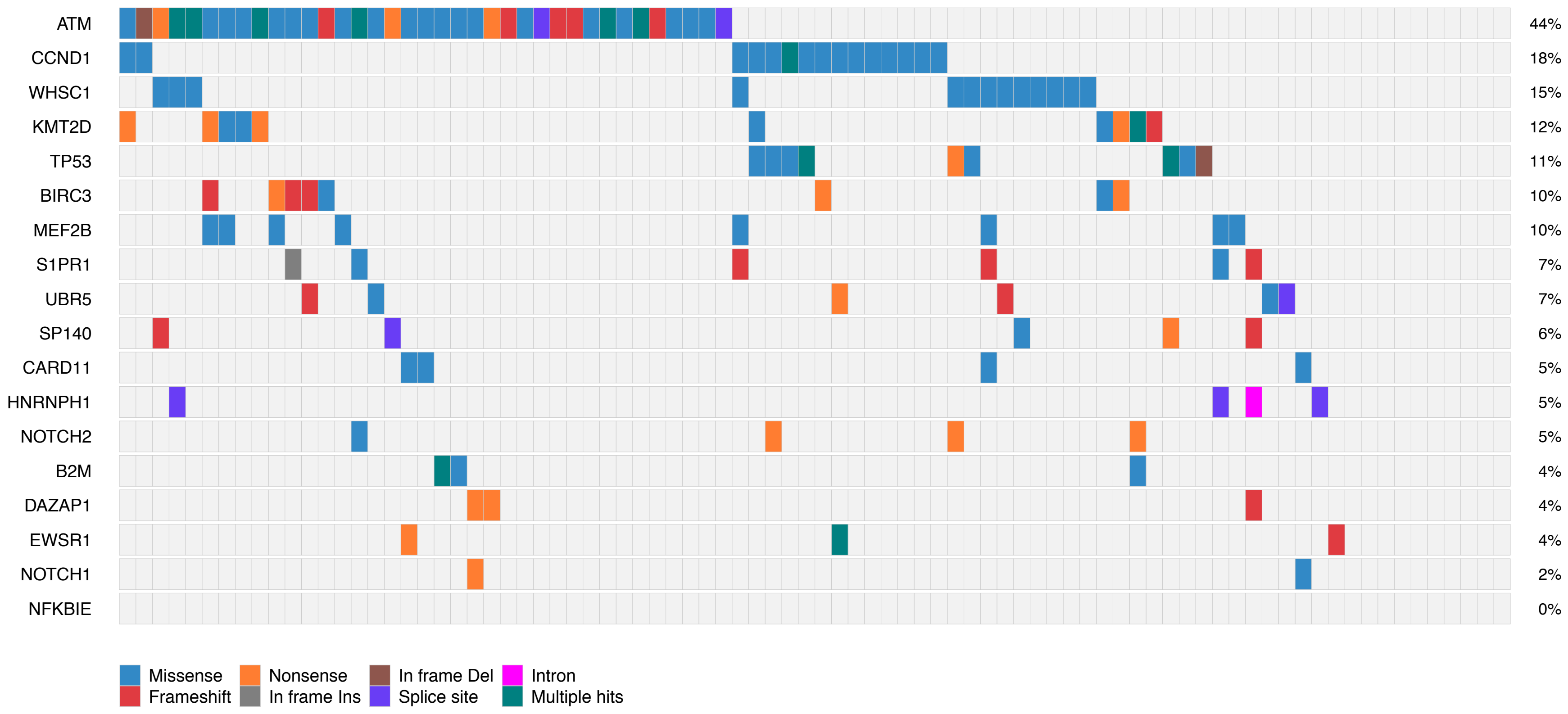


**Supplemental Figure 2. Exome ("Discovery") cohort mutation pattern.** Mutations observed across 84 MCL exome samples in 18 genes used in targeted sequencing. Mutations shown here are limited to non-silent mutations for all genes with the exception of *HNRNPH1*, for which intronic and silent mutations were also included.

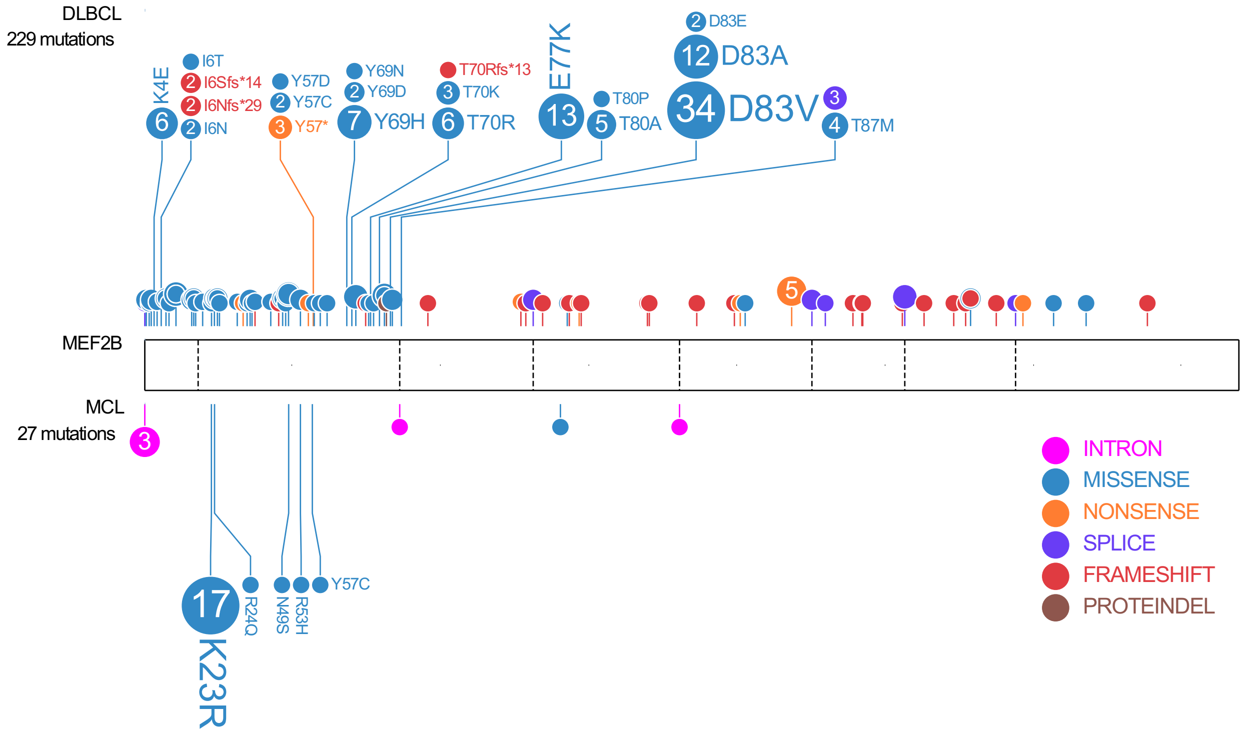


**Supplemental Figure 3. MEF2B Mutation Pattern in MCL and DLBCL**

In both cancers, the majority of mutations affect the MADS/MEF domain with MCLs containing fewer mutations than DLBCLs. *MEF2B* mutations in MCLs predominantly represented K23R whereas DLBCL has more recurrent mutations in this domain and a paucity of K23R.

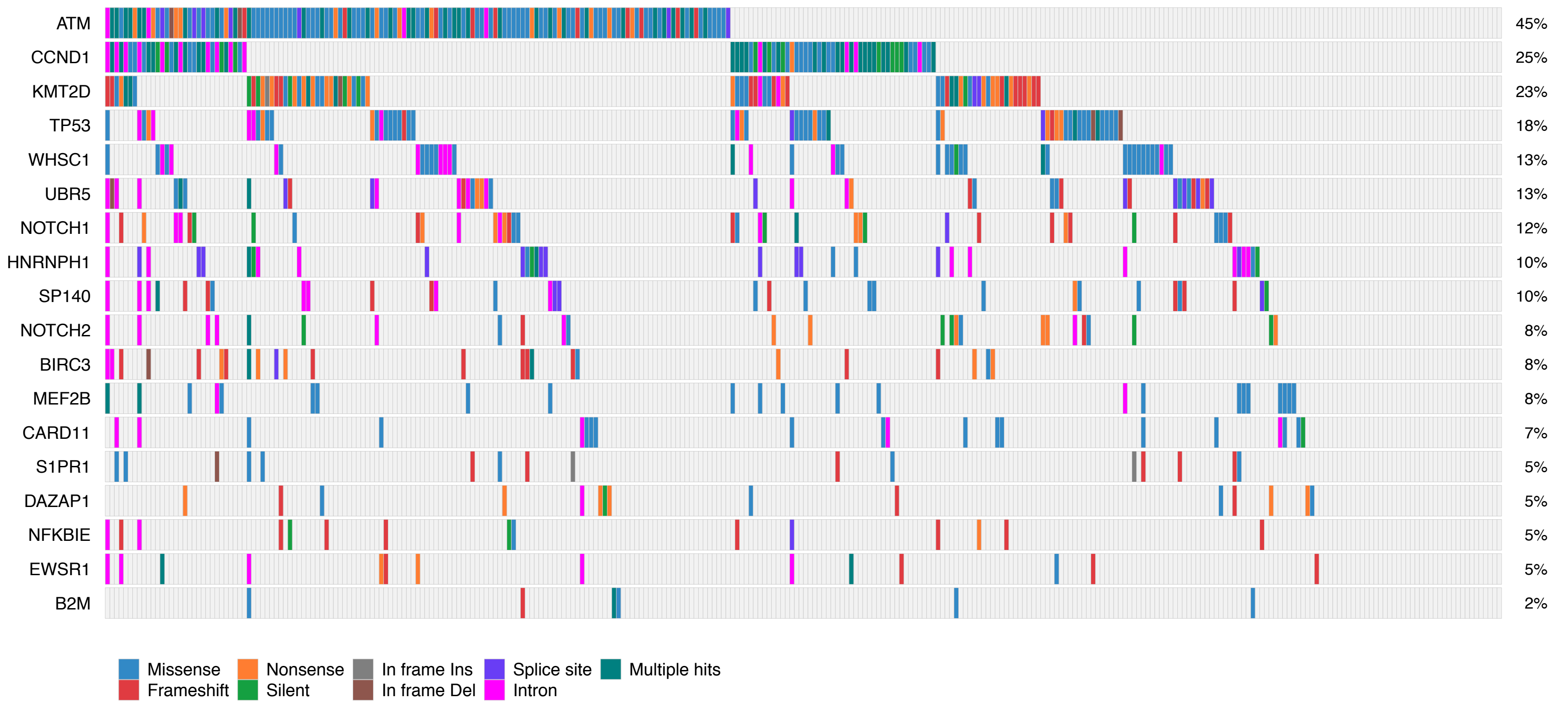


**Supplemental Figure 4. Mutation pattern in MCL including non-coding mutations.** Mutations observed in 291 MCL samples in 18 candidate MCL genes.

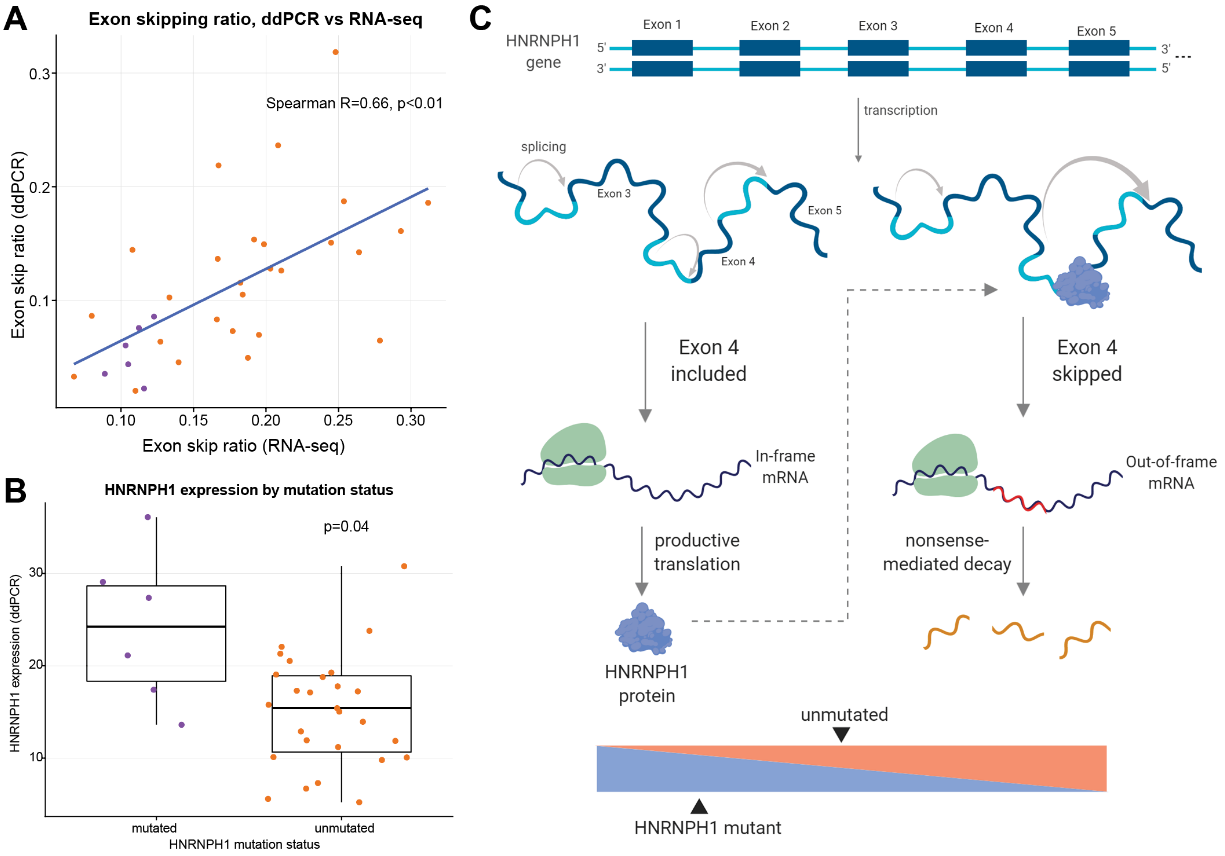


**Supplemental Figure 5. *HNRNPH1* mutations influence expression of HNRNPH1.** (A) Exon-skipping ratios were significantly correlated between RNA-seq and a custom ddPCR assay. (B) In tumors with mutations of *HNRNPH1* (n=6), ddPCR confirmed higher total HNRNPH1 mRNA expression. (C) The proposed model for *HNRNPH1* autoregulation suggests that productive splicing and translation of an mRNA including exon 4 generates HNRNPH1 protein, while the presence of excess HNRNPH1 causes exon 4 skipping and leads to nonsense-mediated decay. The observed mutations proximal to exon 4 disrupt this balance and lead to decreased rates of exon 4 skipping.

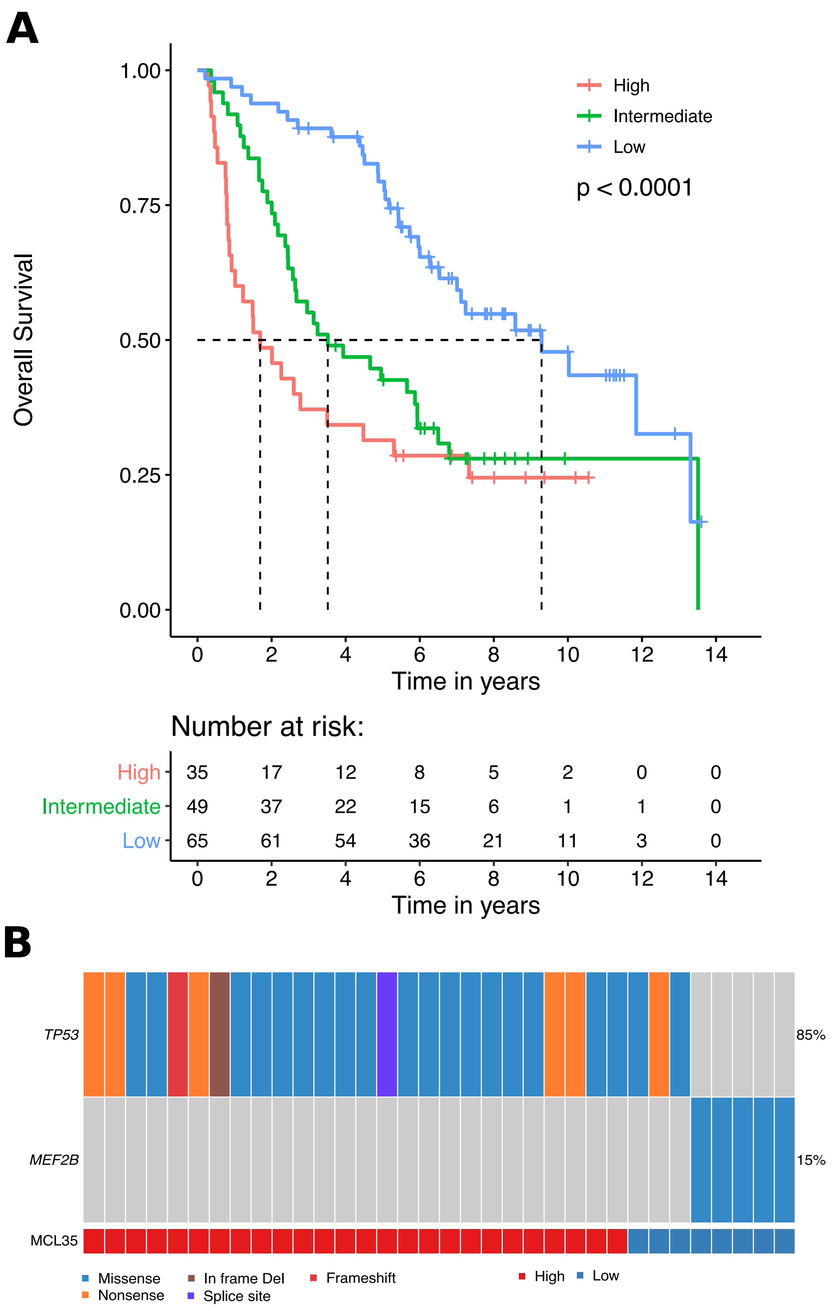


**Supplemental Figure 6. Overall survival (OS) between low-, intermediate-, high-risk MCL35 cases**. (A) Kaplan-Meier method was used to estimate OS and log-rank test was used to assess OS differences between risk groups. Low-risk cases had significantly longer OS relative to both intermediate- and high-risk cases ($P = 5.8 \times 10^{-4}$ and $P = 5.0 \times 10^{-5}$, log-rank test). In contrast, intermediate- and high-risk cases overall survival were not significantly different (P =

1.5 x $10^{-1}$, log-rank test). High and intermediate cases were combined in all relevant analyses and were together considered high-risk cases. (B) *MEF2B* K23R are significantly associated with low-risk cases and *TP53* mutations are associated with high-risk cases.
